# YAMDA: thousandfold speedup of EM-based motif discovery using deep learning libraries and GPU

**DOI:** 10.1101/309872

**Authors:** Daniel Quang, Yuanfang Guan, Stephen C.J. Parker

## Abstract

**Motivation:** Motif discovery in large biopolymer sequence datasets can be computationally demanding, presenting significant challenges for discovery in omics research. MEME, arguably one of the most popular motif discovery software, takes quadratic time with respect to dataset size, leading to excessively long runtimes for large datasets. Therefore, there is a demand for fast programs that can generate results of the same quality as MEME.

**Results:** Here we describe YAMDA, a highly scalable motif discovery software package. It is built on Pytorch, a tensor computation deep learning library with strong GPU acceleration that is highly optimized for tensor operations that are also useful for motifs. YAMDA takes linear time to find motifs as accurately as MEME, completing in seconds or minutes, which translates to speedups over a thousandfold.

**Availability:** YAMDA is freely available on Github (https://github.com/daquang/YAMDA)

## 1 Introduction

*De novo* motif discovery is a common technique for the analysis of biopoylmer sequences such as DNA, RNA, and proteins. It involves the identification of enriched short patterns, commonly referred to as motifs, of monomer letters from a collection of related sequences. One of the most frequent applications of motif discovery is to datasets arising from transcription factor (TF) binding experiments, where motifs correspond to sequence-specific binding patterns of TFs.

MEME (Bailey *et al*., 1994) is a popular probabilistic motif discovery program that uses the expectation-maximization (EM) algorithm to infer motifs as position probability matrices (PPMs), which describe the probability of each possible letter at each position in the pattern. Given a background model, a PPM can be converted to a position weight matrix (PWM) of log odds ratios. MEME uses the batch version of the EM algorithm, which updates parameters after a complete pass through the data. In practice, MEME takes quadratic time relative to the number of letters, leading to prohibitively long run times for large modern high throughput datasets. The majority of the runtime is devoted to seed searching because EM is prone to converging to local optima.

EXTREME is a motif discovery program designed to infer motifs as accurately as MEME in linear time (Quang and Xie, 2014). To achieve this goal, EXTREME uses a word-based discriminative algorithm to search for gapped k-mer words that are enriched in a positive sequence set relative to that of a negative control. Starting points in the search space are derived from the enriched words. Moreover, EXTREME replaces MEME’s batch EM with the online EM algorithm. In contrast to batch learning, online learning updates the parameters after each data sample. Online learning converges faster because it performs multiple parameter updates per data pass instead of one.

Recent advances in “deep learning” offer solutions for improving upon MEME. For example, convolutional neural networks (CNNs) have been shown to be effective for motif discovery (Quang and Xie, 2016). The convolutional layer consists of a set of learnable kernels. The kernels are similar to PWMs, except weights are not constrained to be probabilities or log odds ratios. CNNs are slow to train; however, training can be accelerated through the use of graphics processing units (GPUs) and tensor libraries that are optimized for operations like convolution.

## 2 Software description

YAMDA is a novel program that extends the EXTREME framework by leveraging innovations in deep learning. Specifically, YAMDA uses deep learning libraries to accelerate EM-related computations. Similar to EXTREME, YAMDA’s seeding step uses a discriminative algorithm to find the 100 most enriched gapped k-mer words and converts the words to PWM seeds (see Section 4.4 of Bailey *et al*. (1994)). Initial background probabilities are computed by counting the letter occurrences in the dataset. One “mini-batch” (compromise between batch and online) EM iteration followed by one batch EM iteration is run on each starting point. To parallelize these computations across all seeds, PWMs are treated as convolutional kernels, unloading a bulk of the computational burden on the deep learning libraries and (if available) the GPU. It is for this reason that we chose to use mini-batch EM instead of online EM, since minibatch EM can take advantage of the vectorization. Batch EM is then run to completion on the single seed that yields the highest data likelihood.

## 3 Implementation

YAMDA is built on Pytorch (Paszke *et al*., 2017), a lightweight deep learning Python package with strong support for GPU acceleration; however, YAMDA can also run on the CPU. It accepts FASTA sequences as inputs, and outputs motifs in Minimal MEME format.

## 4 Examples

To demonstrate the efficacy of YAMDA, we use it analyze the 100 bp summit-centered peak repeat-masked sequences from ENCODE TF ChIP-seq datasets, and a digital genomic footprint (DGF) dataset (Quang and Xie, 2014) (Table 1). YAMDA is run in GPU and CPU mode, and both modes are orders of magnitude faster than MEME. Due to MEME’s quadratic runtime, this speedup quickly increases as a function of input size. In comparison, CUDA-MEME (Liu *et al*., 2010), another GPU-accelerated implementation of MEME, demonstrates speedups of less than 1.5, which is orders of magnitude slower than even YAMDA’s CPU mode. These results demonstrate the importance of YAMDA’s linear time seeding; a simple linear speedup of the MEME algorithm is not sufficient since its base runtime grows too fast. Moreover, all of the YAMDA and MEME example output motifs display significant similarity (*E* < 10^−7^) to known motifs in the JASPAR database (Khan *et al*., 2017) according to TOMTOM (Gupta *et al*., 2007). Visually, however, the YAMDA motifs more closely resemble the MEME motifs than the JASPAR motifs, especially for IRF4. This is likely because motif databases are constantly being updated and therefore may not always have the target motif, which explains why the discovered IRF4 motifs aligned to the similar JASPAR IRF1 motif. Together, these results demonstrate how well YAMDA can reproduce MEME’s results in a fraction of the time. Additional details on the implementation and use of YAMDA are given in the Github repository.

**Table 1.**
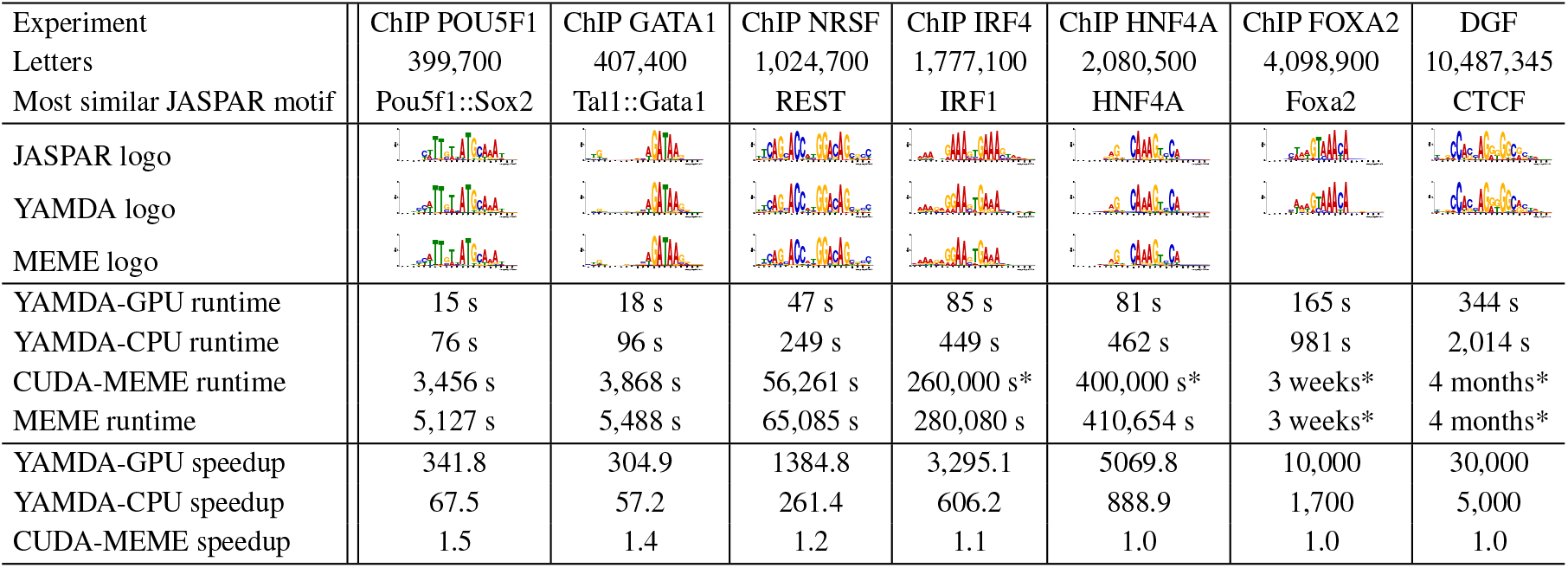
Runtimes for YAMDA (GPU and CPU modes), MEME, and CUDA-MEME to find one motif. Motifs are aligned to the most similar JASPAR motifs. Due to limits in time and resources, some runtimes are estimated. Estimated runtimes are marked with a *.

| Experiment | ChIP POU5F1 | ChIP GATA1 | ChIP NRSF | ChIP IRF4 | ChIP HNF4A | ChIP FOXA2 | DGF |
| --- | --- | --- | --- | --- | --- | --- | --- |
| Letters | 399,700 | 407,400 | 1,024,700 | 1,777,100 | 2,080,500 | 4,098,900 | 10,487,345 |
| Most similar JASPAR motif | Pou5f1::Sox2 | Tal1::Gata1 | REST | IRF1 | HNF4A | Foxa2 | CTCF |
| JASPAR logo |  |  |  |  |  |  |  |
| YAMDA logo |  |  |  |  |  |  |  |
| MEME logo |  |  |  |  |  |  |  |
| YAMDA-GPU runtime | 15 s | 18 s | 47 s | 85 s | 81 s | 165 s | 344 s |
| YAMDA-CPU runtime | 76 s | 96 s | 249 s | 449 s | 462 s | 981 s | 2,014 s |
| CUDA-MEME runtime | 3,456 s | 3,868 s | 56,261 s | 260,000 s* | 400,000 s* | 3 weeks* | 4 months* |
| MEME runtime | 5,127 s | 5,488 s | 65,085 s | 280,080 s | 410,654 s | 3 weeks* | 4 months* |
| YAMDA-GPU speedup | 341.8 | 304.9 | 1384.8 | 3,295.1 | 5069.8 | 10,000 | 30,000 |
| YAMDA-CPU speedup | 67.5 | 57.2 | 261.4 | 606.2 | 888.9 | 1,700 | 5,000 |
| CUDA-MEME speedup | 1.5 | 1.4 | 1.2 | 1.1 | 1.0 | 1.0 | 1.0 |

## Acknowledgements

The NVIDIA Corporation donated the Titan Xp GPU used for development. We thank Vivek Rai for software testing and Tingyang Li for designing the software logo. This work was supported by the National Heart, Lung, and Blood Institute [U01HL137182] to SCJP.

